# MetaPaCS: A Novel Meta-Learning Framework for Pancreatic Cancer Subtype Identification

**DOI:** 10.64898/2025.12.29.696875

**Authors:** Nick B. Peterson, Mengtao Sun, Xinchao Wu, Jieqiong Wang, Shibiao Wan

**Affiliations:** Department of Genetics, Cell Biology and Anatomy, University of Nebraska Medical Center, Omaha, NE; Department of Neurological Sciences, University of Nebraska Medical Center, Omaha, NE; Fred and Pamela Buffett Cancer Center, University of Nebraska Medical Center, Omaha, NE, USA

**Keywords:** Pancreatic cancer, meta learning, Stacking model, Subtype identification, Cancer subtyping

## Abstract

As the third leading cause of cancer related deaths in the United States, pancreatic cancer (PaC) is a highly heterogenous malignancy that can be divided into a multitude of potential subtypes, with the main 4 consisting of aberrantly differentiated endocrine exocrine (ADEX), immunogenic, progenitor, and squamous. Each PaC subtype is characterized by unique molecular pathways and therapeutic characteristics. Identifying PaC molecular subtypes is essential for downstream patient risk stratification and tailored treatment design. Conventional wet-lab approaches for PaC subtyping like microdissection, histopathological studies or molecular profiling are often laborious, costly, and time-consuming. To address these concerns, we present MetaPaCS, a novel meta-learning framework to accurately identify PaC subtypes based on transcriptomics data only. Specifically, after preprocessing, the transcriptome-based feature vectors were classified by 10 base machine learning (ML) classifiers, whose prediction outputs were then combined with the initial preprocessed feature vectors to constitute a new set of ensemble feature vectors for a meta-learning model. Our meta-learning model could learn and leverage the diversity of different base classifiers to boost the prediction performance beyond any single ML model. Results based on 100 times ten-fold cross validation tests on benchmarking datasets demonstrated that MetaPaCS performed significantly better than existing state-of-the-art methods for PaC subtyping. In addition, our meta-learning model remarkably outperformed each individual base classifier, demonstrating that MetaPaCS could combine diverse results from multiple base classifiers to boost the ensemble performance. We believe that MetaPaCS is a promising tool for characterizing PaC subtypes and will have positive impacts on downstream risk stratification and personalized treatment design for PaC patients.

## Introduction

Pancreatic cancer (PaC), a highly lethal malignancy that is nearly undetectable in its earlier stages, is the 3^rd^ leading cause of death in the USA, and is projected to become the 2^nd^ within a decade^1,2^. Since 1970, the amount of new PaC cases per year has increased drastically from 18,800 to 64,050 in 2023^3^. In contrast, PaC’s 5 year survival rate has only improved by 3% in that same time frame, going from a 9% 5 year survival rate to a 12% 5-year survival rate while breast cancer’s 5 year survival rate for has improved from 40-50% in 1970 to 90-100% in 2023 in that same time frame^4^. Additionally, PaC’s mutational landscape is highly heterogenous among patients and within the tumor itself, making it difficult to find a singular effective treatment for all PaC cases. While current more generalized treatments like neoadjuvant therapy have resulted in slight improvements, the most promising improvements have instead come from experimental targeted treatments such as immunotherapy using novel delivery methods^5^. The effectiveness of these new personalized treatment options highlights the benefits of researching the variation that can be found in PaC’s different molecular pathways.

Subtyping has become an increasingly popular way to provide more personalized healthcare. A notable example is the improved prognostic and predictive performance for breast cancer^6^ when the subtype information is incorporated into diagnosis. An initial study in 2015 found 2 distinct subtypes of PaC^7^, namely basal and classical, though even within these subtypes there is high variation between each individual case. Further PaC subtyping^1^ has been conducted via expression analysis correlating with previously seen histopathological characteristics from microdissection, identifying 4 key subtypes of PaC: aberrantly differentiated endocrine exocrine (ADEX), immunogenic, progenitor, and squamous. Each subtype shows differing characteristics in treatment and survival^1^. Specifically, the immunogenic subtype is associated with upregulation in genes involved with B cell signaling pathways, antigen presentation, CD4+ T cell signaling, CD8+ T cell signaling, and Toll-like receptor signaling pathways, creating an immunosuppressant microenvironment. As a result, this subtype would likely respond less to immunotherapy. The squamous subtype shows mutations similar to those seen in other C2-squamous-like class tumors of different cancers, and may share therapeutic strategies. These unique characteristics ultimately make subtyping each PaC case a logical choice for both creating the most efficient individual treatment plans and for minimizing the side effects of the more general treatments.

Conventionally, wet-lab based approaches are used for identifying PaC subtypes, with the subtypes defined histopathologically being found through microdissection and examination of the samples under a microscope^7^, which is labor intensive, requires the resection of the tumor, and may be infeasible for the patients where tumor resection is not possible. Biomarker-based methods are becoming another increasingly used type of wet-lab approach^8^ for subtyping for other cancers^9^, though they are currently not suitable for PaC subtyping. Ca19-9^3^, the only FDA approved biomarker for PaC, has a low sensitivity and often leads to late diagnoses, which is not suitable for accurate PaC subtyping. In addition, Ca19-9 is ineffective for diagnosis in the 5 to 10% of people who possess the Lewis negative phenotype due to the reduced synthesis of Ca19-9^10^. While other promising biomarkers are currently being researched for diagnosis such as MIC, TNP1, and RIT2^3^, they are still in the early stages of testing and are unable to differentiate between subtypes. In summary, current wet-lab methods to characterize PaC subtyping are time-consuming, labor-intensive, expensive, or even ineffective.

Computational methods offer an alternative solution for efficient and cost-effective PaC subtyping, as the omics data of a tumor is much less invasive to obtain and does not require the presence of large tumors for analysis. Computational methods like machine learning approaches have been widely applied to different aspects of cancer subtyping studies, including risk stratification prediction^1112^, subtype characterization^13,14^, and subtype prediction^15^. Due to their versatility and ability to handle large amounts of data, computational methods can make full use of features that may otherwise be difficult for a human researcher, including clinical data, omics data^16^, image data^17^, or multi-modal data. Specific methods for PaC subtype prediction include the use of nonnegative matrix factorization (NMF) derived meta-genes in H2O^18^. Another study^19^ makes use of multiple deep learning models, including Vanilla, IDaRS, DeepMIL, and VarMIL, to predict PaC subtype based on whole slide pathological images.

While these machine learning approaches have demonstrated different levels of competitiveness for PaC subtyping, they may not perform robustly and consistently across different datasets and cases due to limited generalizability. Stacking is an ensemble learning technique which uses multiple different classifiers in its prediction, preventing the model from overfitting and reducing the overall influence of bias, thus enhancing the model’s generalization capabilities^20^. A stacking model makes use of multiple base classifiers with different strengths and characteristics to generate a set of initial prediction probabilities which can then be used as input data for a meta-learning classifier. This allows the meta-learning classifier to learn from the base classifier’s biases and account for them, improving the robustness of the results^21^. Similar models have shown high performance for other biological tasks like Alzheimer’s diagnosis^22^, lung cancer characterization^23^, and breast cancer in past studies^20^.

To this end we propose a novel meta-learning approach, MetaPaCS, for the accurate and reliable prediction of PaC subtypes based on transcriptomics data only. Transcriptomics data based machine learning models have been successfully used in the subtyping of multiple other cancers, such as B-cell acute lymphoblastic leukemia^24–26^, ovarian cancer^27^, and medulloblastoma^26^ due to its ability to provide a direct look at molecular pathway activity within a tumor. MetaPaCS makes use of multiple base classifier algorithms in a stacking model for both a more robust classification scheme and a reduced risk of algorithmic bias. We demonstrate that MetaPaCS can both outperform any individual machine learning classification method in subtype classification tasks and is shown to improve the visualization of PaC subtypes when compared to methods like PCA, UMAP, and t-SNE.

## Methods

### Dataset collection and preprocessing

In this study, we used the RNA-seq expression data of 96 PaC patients from ^1^. These samples, all between 34 and 90 years of age, are classified into four different subtypes (**Fig. 1A-B**). To prevent the possible skewing of results, missing values in the expression data were replaced with the average expression value for that same gene across all samples using imputation. The data was converted back into counts per million (CPM) before the coefficient of variation squared (CV^2^) was calculated for each gene. Only genes with a CPM above 1 and a CV^2^ above 0.5 were kept, reducing the number of genes from 18,276 to 368 and preserving only the most varied genes for classification (**Fig. 1C**).

**Fig. 1.**
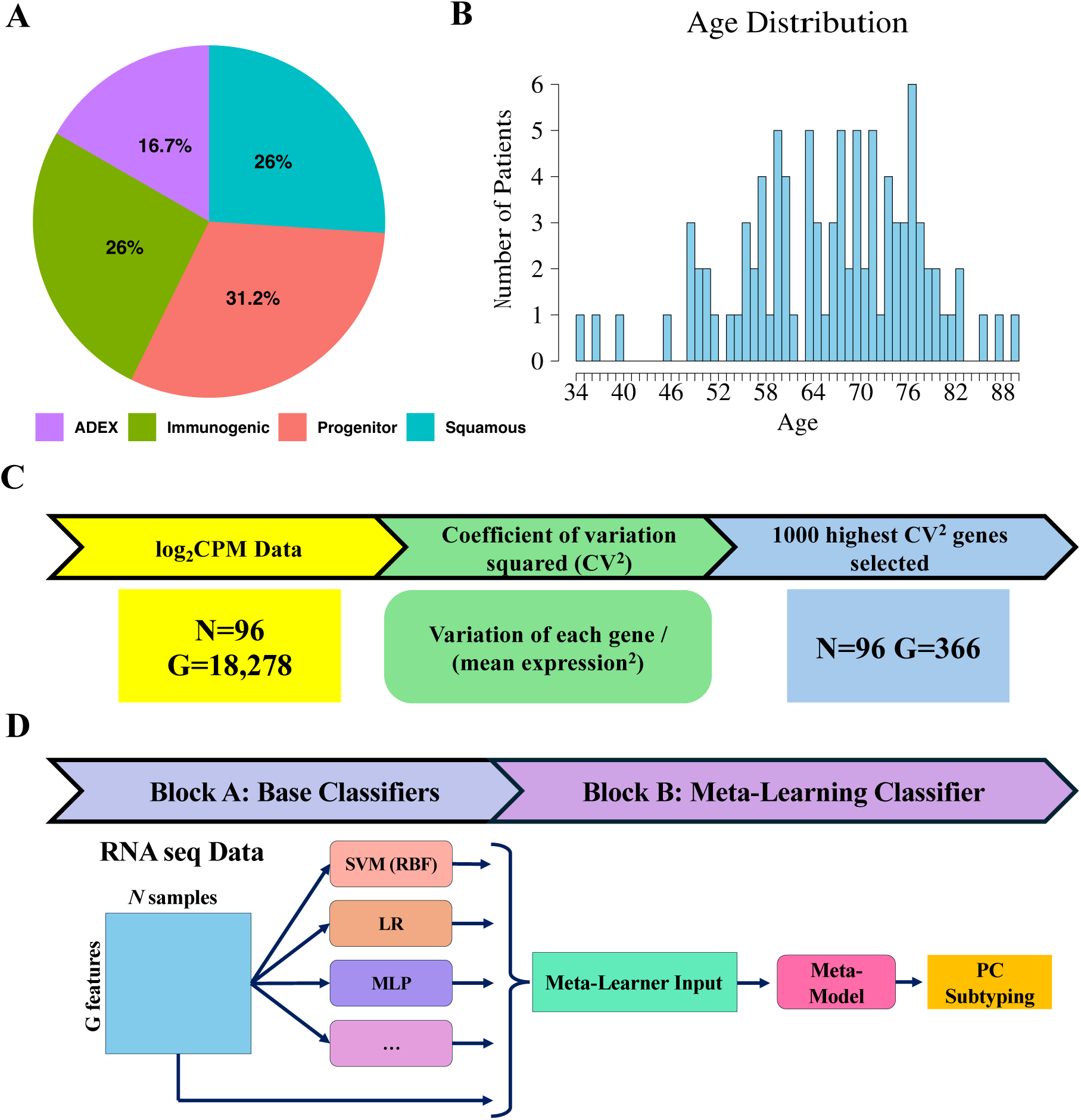
Overview of the MetaPaCS framework. (**A**) Distribution of subtypes within the dataset. (**B**) Age demographics of the patient cohort. (**C**) Initial expression data undergoes log transformation followed by empirical Bayes moderation to identify significantly different genes between subtypes, thereby reducing dimensionality. (**D**) Stacking model architecture: processed data is input into ten distinct base classifiers (SVM (linear), SVM (RBF), XGB, RF, KNN, LR, NB, DT, QDA, and MLP). The resulting predictions are then concatenated with the preprocessed input data and fed into the meta-learner (SVM (linear)).

### Meta learning for pancreatic cancer subtyping

For PaC subtyping, we propose MetaPaCS, a novel meta-learning framework for accurate identification of PaC subtypes based on transcriptomic data only. Our model is made up of 2 main blocks, block A (the base classifier algorithms), and block B (the meta-learning classifier algorithm) (**Fig. 1D**). Block A uses the filtered expression data as input for 10 different base classifiers to produce an ensemble output containing the subtype probability predictions for each base classifier. The output would then be concatenated to the original input data, serving as input for Block B’s meta learning classifier to create a final subtype probability prediction.

There are a total of 10 different base classifiers used in Block A: Support vector machines (SVMs) with either a linear kernel (SVM linear)^28^ or a radial basis function kernel (SVM RBF)^29^, Decision tree classifier (DT)^30^, random forest classifier (RF)^31^, XGBoost (XGB)^32^, Gaussian naïve bayes (NB)^33^, Logistic regression (LR)^34^, Multilayer perceptron (MLP)^35^, K-nearest neighbors (KNN)^36^, and Quadratic discriminant analysis (QDA)^37^. These classifiers were chosen due to their diverse methods of analysis, ensuring a variety of predictions for a more holistic predictive performance^38^. To better enhance the performance of the base classifiers and quality of predictions, some of the possible classifiers chosen were ensemble methods themselves.

MetaPaCS next concatenates the probability outputs generated by the Block A base classifiers with the original filtered expression matrix, producing an augmented feature space that integrates both molecular expression profiles and model-derived predictive signals. This combined dataset is subsequently used as input to Block B, the meta-learning stage, where an independent classifier is trained to generate the final prediction by explicitly learning from the complementary strengths and algorithmic biases of the base models. A linear support vector machine (SVM) was selected as the primary meta-classifier. To determine the optimal meta-learning strategy, six candidate classifiers were systematically evaluated, including linear SVM, logistic regression (LR), multilayer perceptron (MLP), decision tree (DT), and random forest (RF). Model performance was assessed using prediction accuracy, and the best-performing classifier was adopted for the final MetaPaCS framework. Detailed hyperparameters for all classifiers used in Blocks A and B are provided in **Supplemental Table S1**.

Leave-one-out cross-validation (LOOCV) was used to evaluate the performance of MetaPaCS and other state-of-the-art approaches for PaC subtyping. Nine different metrics were used to provide a more comprehensive evaluation of a model’s effectiveness: accuracy, area under curve (AUC), Matthews Correlation Coefficient (MCC), F1-Score, recall, G-measure, specificity, Jaccard index, and precision.

### Model testing and optimization

To optimize our framework, we tested multiple combinations of base and meta-learning classifiers to better analyze the learning process and find the highest performing classifier combinations for each block. First, different numbers of base classifiers were tested to see if the additional prediction data would improve the meta-learning performance. Block A models including 2, 4, 6, 8, and 10 of the most accurate base classifiers were evaluated for all 6 of the possible meta-learning classifiers in block B for comparison (**Supplemental Fig. S1**). Additionally, we tested the effects of different classifier combinations on the meta-learning performance of the model by evaluating the model’s performance on all unique combinations of 2 classifiers from the 7 best performing base classifiers in block A with the model performance being evaluated for all 6 possible meta-learning classifiers in block B (**Supplemental Fig. S2**).

Other additional classifiers were used due to factors such as relatively low computational burden and good performance. KNN, LR, NB, and DT were chosen due to their high computational efficiency^39,40^. QDA and MLP were chosen due to their relatively high performance, with QDA being highly effective on classes with different covariances and MLP being able to iteratively learn data patterns with backpropagation^41^. Both linear and radial basis function (RBF) SVM’s were used for their strong performance in high dimensional data, which is especially useful in the high-dimensional RNA seq data.

### MetaPaCS Visualization

To capture both the global structure of the data and the specificity of subtype predictions for MetaPaCS,, we made use of UMAP, t-SNE, and PCA using prcomp, the Rtsne, and uwot packages to evaluate the effectiveness of MetaPaCS at visualization compared to basic visualization methods. Specifically, MetaPaCS utilized a combination of two key matrices: a subtype score matrix and a sample-to-subtype matrix derived from prediction results. The former matrix was obtained through the MetaPaCS subtype identification workflow (**Fig. 1**), in which subtype-specific scores were calculated for each sample. This matrix captured the continuous prediction confidence of each sample across the four PaC subtypes, forming a *S* × *L* matrix, where *S* is the number of samples and *L* is the number of subtypes. A sample-to-subtype matrix was constructed using one-hot encoding based on the predicted subtype labels derived from the subtype score matrix. Specifically, the subtype with the highest score for a given sample was assigned a value of 1, while all other subtypes were assigned 0. This results in a binary matrix of the same dimension (*S* × *L*), representing discrete subtype assignments. These two matrices were then concatenated along the feature axis to form a unified visualization matrix, which was used as input for t-SNE, PCA and UMAP to generate a low-dimensional embedding. This integration strategy enabled the combination of both quantitative prediction certainty and categorical class identity, thereby enhancing the biological interpretability and subtype-level separation in the visualization space.

### Subtype-specific differential gene expression analysis

Subtype-specific differential gene expression analysis was conducted via one-vs-rest comparisons for each subtype. Genes were considered subtype specific if they meet criteria of *p* value < 0.05 and log2 Fold Change > 1. The top 10 most variable genes from each subtype were then visualized in a heatmap to highlight the transcriptional difference across subtypes. Additionally, volcano plots were generated for each subtype with a threshold of |log2FC| > 1.5, and adjusted *p* value < 0.05. Finally, t-SNE plots were generated based on differential expression profiles, with samples colored by the expression of a significantly up-regulated gene corresponding to a specific subtype.

### Gene Ontology (GO) Pathway analysis

The top 500 up-regulated genes for each subtype were subjected in Gene Ontology (GO) using ClusterProfiler^42^ (V3.21) enrichment analysis to identify the top 15 most significantly enriched pathways within Biological Processes, Cellular Components, and Molecular Function. Gene-Drug-Disease Association Studies

### Gene-Drug-Disease Association Studies

Drug and disease linkages were extracted from the DrugBank database^43^ and the target-disease linkages were curated in the Therapeutic Target Database (TTD)^44^. The up and downregulated genes were cross-referenced against these pharmacological datasets to give a list of drugs for both possible novel therapeutic agents for each cancer subtype and potential effects each subtype may have on the metabolism and function of these existing agents.

## Results

### Study design for PaC identification

To accurately identify PaC subtypes using transcriptomic data, we developed MetaPaCS, a meta-learning based framework designed to integrate heterogeneous machine-learning signals for improved subtype prediction. Based on 96 collected PaC patients (**Fig. 1A–B**), their gene expression profiles were reconstructed in CPM format, followed by gene filtering using CV² to retain the most informative features. These processed data served as input for the two-block architecture of MetaPaCS (**Fig. 1C–D**). Block A generated an ensemble of subtype probability predictions by applying a diverse set of base classifiers, each capturing different aspects of the high-dimensional transcriptomic landscape. The outputs from Block A were then concatenated with the original filtered gene expression data, forming an enriched representation that was subsequently used by block B, the meta-learning module. Block B integrated these complementary predictive signals to refine decision boundaries and produced the final subtype prediction. To optimize MetaPaCS, different combinations of base learners and meta-learners across multiple configurations were systematically compared. Incorporating the full set of base classifiers in block A and using a linear SVM as the meta-learner in Block B resulted in the most consistent and accurate subtype predictions.

### Model optimization and benchmarking for identifying PaC subtypes

To determine the optimal architecture of MetaPaCS, a wide range of classifier combinations for Block A and Block B were systematically evaluated. Each of the ten base classifiers in block A were first trained individually and ranked according to performance to provide a reference for model selection. Subsequently, the influence of different base-classifier combinations on meta-learning performance was assessed. Block A was constructed using the top 1, 2, 4, 6, 8, or all 10 classifiers, and each configuration was exhaustively paired with six candidate meta-learners in block B (**Supplemental Fig. S1**). Additionally, all possible pairs of 2 base classifiers were tested from the 7 highest performing base classifiers to evaluate the impact of each individual base classifier on the performance of block B. This comprehensive search enabled evaluation of how classifier diversity and ensemble breadth affected overall predictive performance (**Supplemental Fig. S2**). Across all tested configurations, model performance either remained stable or improved as additional base classifiers were included in block A. Linear SVM consistently emerged as the strongest performing meta-learner in block B when all 10 base classifiers were used (**Supplemental Fig. S3**). Based on these results, the optimal MetaPaCS architecture consisted of all ten base classifiers in Block A and a linear SVM as the block B meta-learner, which provided the most robust and accurate subtype predictions across accuracy, precision, recall, F1-score, MCC, G-measure, AUC, specificity and Jaccard index.

The predictive performance of the optimized MetaPaCS framework was benchmarked against each of the ten independently trained base classifiers (**Fig. 2**). These individual classifiers were first trained and performance-ranked to establish a comparative reference. Across most evaluated metrics, including accuracy, precision, recall, F1-score, MCC, G-measure, specificity, and Jaccard index, MetaPaCS consistently achieved the highest performance, outperforming all single-model classifiers in all performance metrics (**Fig. 2A, C-I**) except AUC. For the AUC metric (**Fig. 2B**), MetaPaCS ranked second overall, exhibiting performance 0.4% slightly below that of the top-performing logistic regression model. Despite this exception, MetaPaCS demonstrated the most balanced and robust predictive profile across the full set of evaluation measures. These findings highlighted the advantage of the meta-learning architecture, which could integrate complementary information from diverse base learners to produce more stable and accurate subtype predictions compared with individual classifiers.

**Fig. 2.**
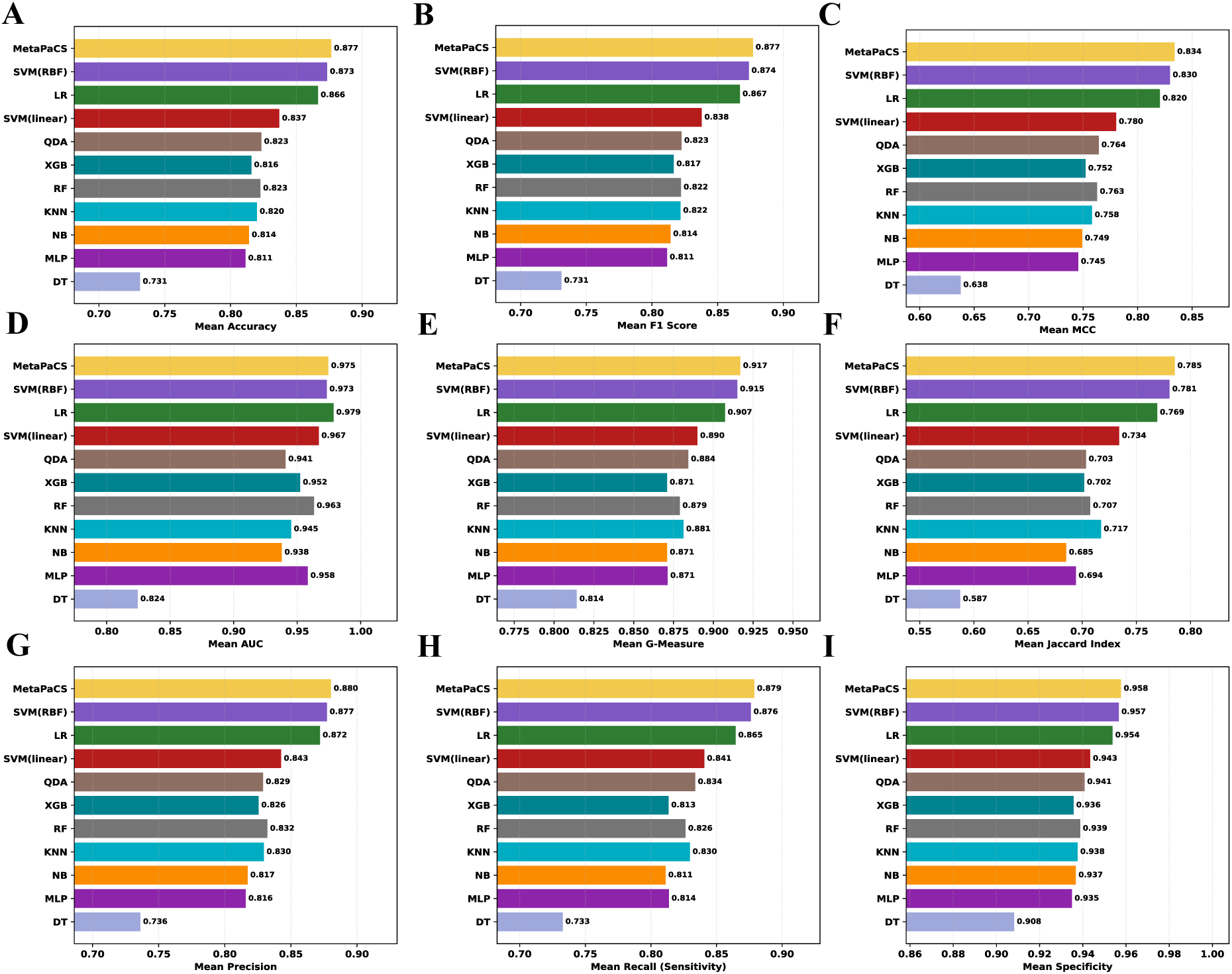
Performance comparison between all base classifiers and MetaPaCS. The y-axis denotes the individual base classifiers and MetaPaCS, while the x-axis represents the value for each evaluation metric. Subfigures display results for: (**A**) Accuracy, (**B**) Area Under the Curve (AUC), (**C**) F1 Score, (**D**) Matthews Correlation Coefficient (MCC), (**E**) Recall, (**F**) Precision, (**G**) Specificity, (**H**) G-Measure, and (**I**) Jaccard Index.

### Subtype-specific differential gene expression analysis showed distinct patterns of expression separating the PaC subtypes

To assess the impact of MetaPaCS on visualizing pancreatic cancer subtypes, three commonly used dimensionality-reduction and visualization methods, including Principal Component Analysis (PCA), t-distributed Stochastic Neighbor Embedding (t-SNE), and Uniform Manifold Approximation and Projection (UMAP), were applied to different feature representations (**Fig. 3**). When these methods were applied directly to the raw gene-expression matrix (**Fig. 3A-C**), samples from the four subtypes (Squamous, ADEX, Progenitor, and Immunogenic) showed extensive overlap, and clear subtype boundaries were not apparent. This pattern indicates that conventional unsupervised visualizations based solely on expression profiles are insufficient to resolve the underlying subtype structure. In contrast, substantially improved clustering was observed when the same dimensionality reduction methods were applied to the MetaPaCS derived embedding (**Fig. 3D-F**), where samples formed more compact and well-separated clusters, reflecting that MetaPaCS captures discriminative structure that was not evident in the original expression space. Visualization was further enhanced when the MetaPaCS embedding was concatenated with one-hot encoded subtype probabilities before dimensionality reduction (**Fig. 3G–I**), resulting in tightly clustered groups with minimal inter-subtype overlap. Among the three visualization methods, t-SNE applied to the MetaPaCS-based representation produced the most distinct separation of subtype clusters, consistent with its emphasis on preserving local neighborhood structure. These findings indicate that MetaPaCS provides a more informative, subtype-aware feature space that markedly improves the effectiveness of standard visualization techniques, with t-SNE being selected as the primary method for subsequent graphical representation of subtype structure.

**Fig. 3.**
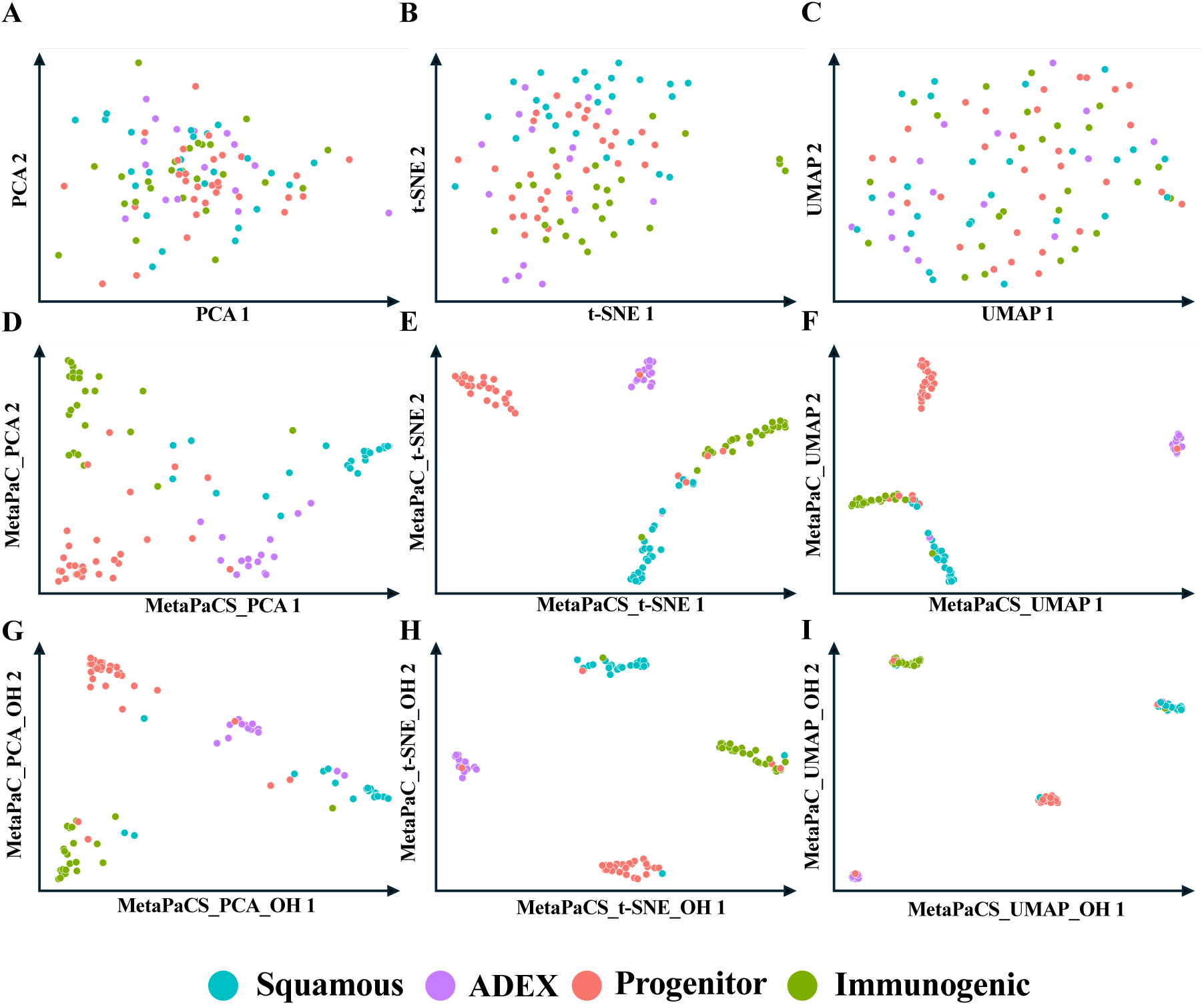
Comparative analysis of dimensionality reduction techniques on filtered gene expression data and corresponding subtype prediction probabilities. Each point represents a sample, colored according to its true subtype. Rows illustrate different strategies: (**A, B, C**) the original filtered data displayed with PCA, t-SNE, and UMAP, (**D, E, F**) subtype prediction probabilities from MetaPaCS displayed with PCA (MetaPaCS_PCA), t-SNE (MetaPaCS_t-SNE), and UMAP (MetaPaCS_UMAP), and (**G, H, I**) subtype prediction probabilities from MetaPaCS concatenated with one-hot encoded (OH) results displayed with PCA (MetaPaCS_PCA_OH), t-SNE (MetaPaCS_t-SNE_OH), and UMAP (MetaPaCS_UMAP_OH). Columns represent the dimensionality reduction technique applied: Principal Component Analysis (PCA; V1) (**A, D, G**), t-distributed Stochastic Neighbor Embedding (t-SNE; V2) (**B, E, H**), and Uniform Manifold Approximation and Projection (UMAP; V3) (**C, F, I**).

### Subtype-specific differential gene expression analysis showed distinct patterns of expression and potential biomarkers for PaC subtypes

To delineate subtype-specific expression patterns and potential biomarkers for PaC, differential gene expression analysis was performed (**Fig. 4**). As shown in the heatmap and volcano plots (**Fig. 4A-E**), the four subtypes exhibited distinct signatures, with sets of genes selectively and significantly up- or downregulated in each subtype. These subtype-specific patterns highlighted underlying biological differences and revealed potential biomarkers associated with their clinical characteristics. Progenitor displayed significant upregulation of G6PC2, a member of the glucose-6-phosphatase catalytic subunit family that participates in the final steps of gluconeogenic and glycogenolytic pathways. Although G6PC2 lacks phosphohydrolase activity, it is selectively expressed in pancreatic islets and plays a critical role in modulating basal glucose homeostasis through the regulation of glucose-6-phosphate handling. Dysregulation of G6PC2 has been associated with alterations in fasting blood glucose levels and pancreatic β-cell function, suggesting that its elevated expression in the Progenitor subtype may reflect subtype-specific metabolic reprogramming consistent with pancreatic lineage features^45^. Immunogenic showed a high amount of upregulated immunoglobin genes such as IGKV1.39 and LGKV6D.21^46^. The Squamous subtype exhibited elevated expression of epithelial and migration-related markers, including MIR205HG and KRT6C. MIR205HG, the host transcript of miR-205, has been linked to epithelial differentiation and enhanced invasive potential in squamous-associated malignancies. KRT6C, a cytokeratin enriched in proliferative squamous epithelia, has been shown to drive tumor cell proliferation, migration, and EMT activation, and its high expression correlates with poor prognosis and aggressive behavior in epithelial cancers such as lung adenocarcinoma^47,48^. The ADEX subtype exhibited marked upregulation of pancreas-enriched digestive enzymes, including CLPS and CELA3B. CLPS encodes colipase, an essential cofactor for pancreatic lipase and a key component of the exocrine digestive system, reflecting strong pancreatic lineage identity. CELA3B, which is exclusively expressed in the exocrine pancreas, encodes a chymotrypsin-like elastase whose dysregulation has been mechanistically linked to chronic pancreatitis, β-cell dysfunction and increased susceptibility to pancreatic adenocarcinoma as demonstrated in both human genetic studies and CRISPR-engineered mouse models^49^. Additionally, the expression level of upregulated DEGs, including *CELA3B*, *PCG*, *NPSR1* and *GPR87*, across all 96 samples highlighted their specific overexpression in ADEX, immunogenic, progenitor and squamous respectively (**Fig. 4F-I**).

**Fig. 4.**
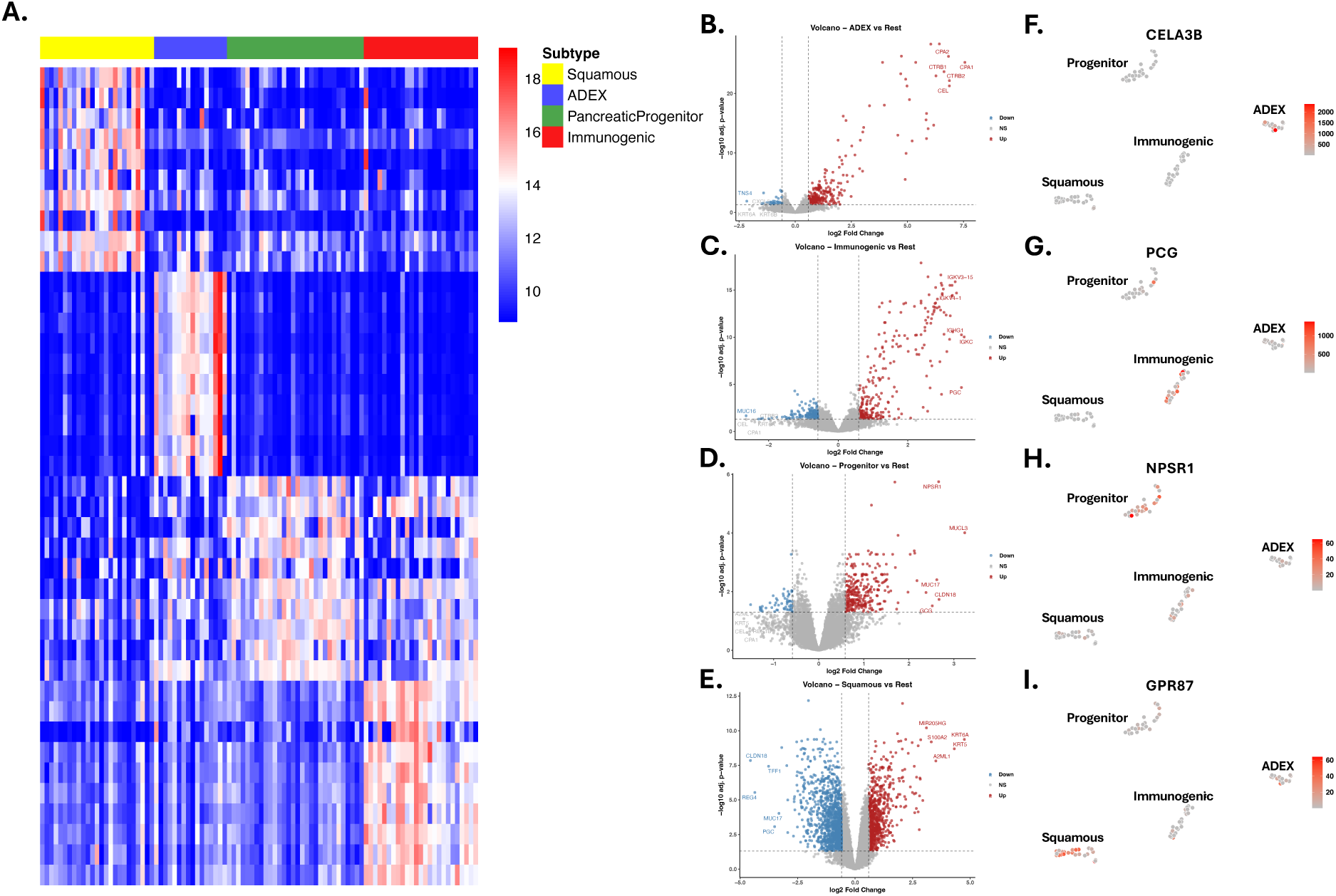
Subtype-specific differential gene expression (DGE) analysis and relative expression visualization. (**A**) Heatmap illustrating the top ten upregulated genes for each subtype in a one-vs-rest comparison. (B-E) Volcano plots highlighting significantly up- and downregulated genes (Log Fold Change [logFC] > 1.5 or < -1.5, and p-value < 0.05) for: (**B**) ADEX, (**C**) Immunogenic, (**D**) Progenitor, and (**E**) Squamous subtypes. The five genes with the highest and lowest logFC values are labeled. (**F-I**) t-SNE plots showing the relative expression (red intensity) of a key upregulated gene within each respective subtype: (**F**) ADEX, (**G**) Immunogenic, (**H**) Progenitor, and (**I**) Squamous.

### Subtype-specific pathway analysis identified distinct enriched pathways for PaC subtypes

The difference between subtypes was further examined through the 15 most prevalent GO pathways of the top 500 upregulated genes in each (**Fig. 5**). The progenitor subtype (**Fig. 5A**) mainly displayed upregulated cellular transport and secretion associated genes, with the most notable upregulated gene being insulin secretion. The squamous subtype (**Fig. 5B**) showed a high upregulation in both internal and external cellular structure related pathways, such as epidermis development and intermediate filament organization. Additionally, 4 of the 15 upregulated pathways in the squamous subtype are involved in kidney cell differentiation. The immunogenic subtype’s (**Fig. 5C**) most upregulated pathways were nearly all immunity pathways, with the only exception being the 17 genes involved in the digestion pathway. The ADEX subtype’s (**Fig. 5D**) most upregulated pathways were mostly based on the secretion of different peptides and hormones, with only 3 pathways based on transport and 2 based on localization.

**Fig. 5.**
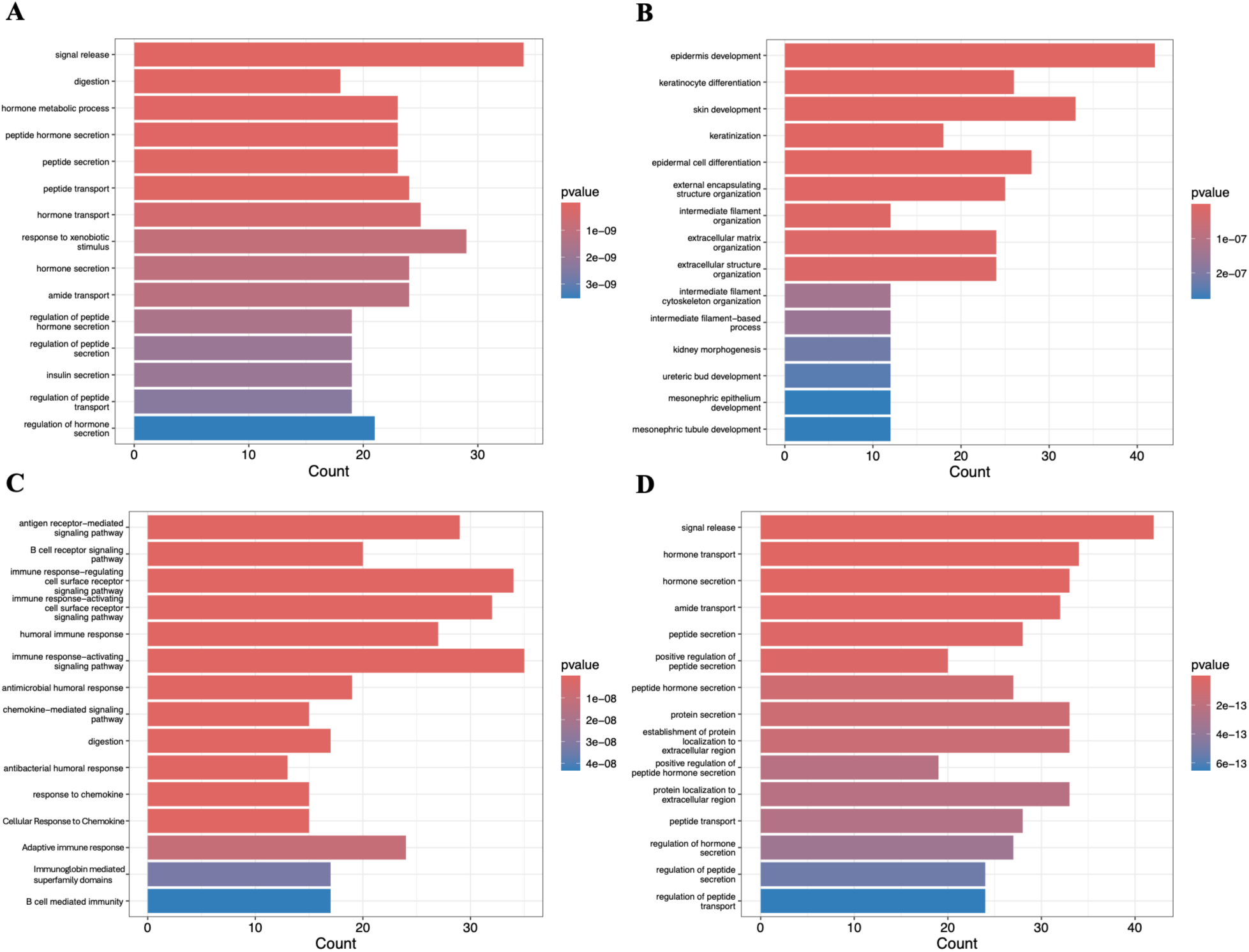
Prominent Gene Ontology (GO) pathways associated with upregulated genes in each subtype. Bar plots illustrating the 15 most significant GO pathways enriched among the top 500 upregulated genes for: (**A**) Progenitor, (**B**) Squamous, (**C**) Immunogenic, and (**D**) ADEX subtypes.

### Drug-disease association studies identified potential subtype-specific target genes and drugs

Pathway analysis was then conducted on these differentially expressed genes for associated drugs and diseases using the drugbank database (**Fig. 6**). We found many of these subtype specific upregulated genes were linked to a multitude of drugs and diseases. In the progenitor subtype (**Fig. 6A**) for example, the upregulated gene NPSR1 was linked to the metabolism of halothane, a commonly used anesthetic in developing countries that is known for severe hepatotoxic side effects^50^. The squamous subtype (**Fig. 6B**) showed upregulation in the gene SLCO1B3, a gene that interacts with gemfibrozil, a drug responsible for preventing high cholesterol/fat levels in the blood^51^. TNFSRF17, which is associated with the metabolism of 5 different medicines that are used for the treatment of multiple myeloma, was found to be upregulated in the immunogenic subtype (**Fig. 6C**). The upregulated genes of the ADEX subtype (**Fig. 6D**) show interactions with 15 different drugs, with several being associated with diabetes, obesity, and other forms of non-pancreatic cancer. Additionally, the gene AMY2A, responsible for starch digestion, was notably upregulated in the ADEX subtype.

**Fig. 6.**
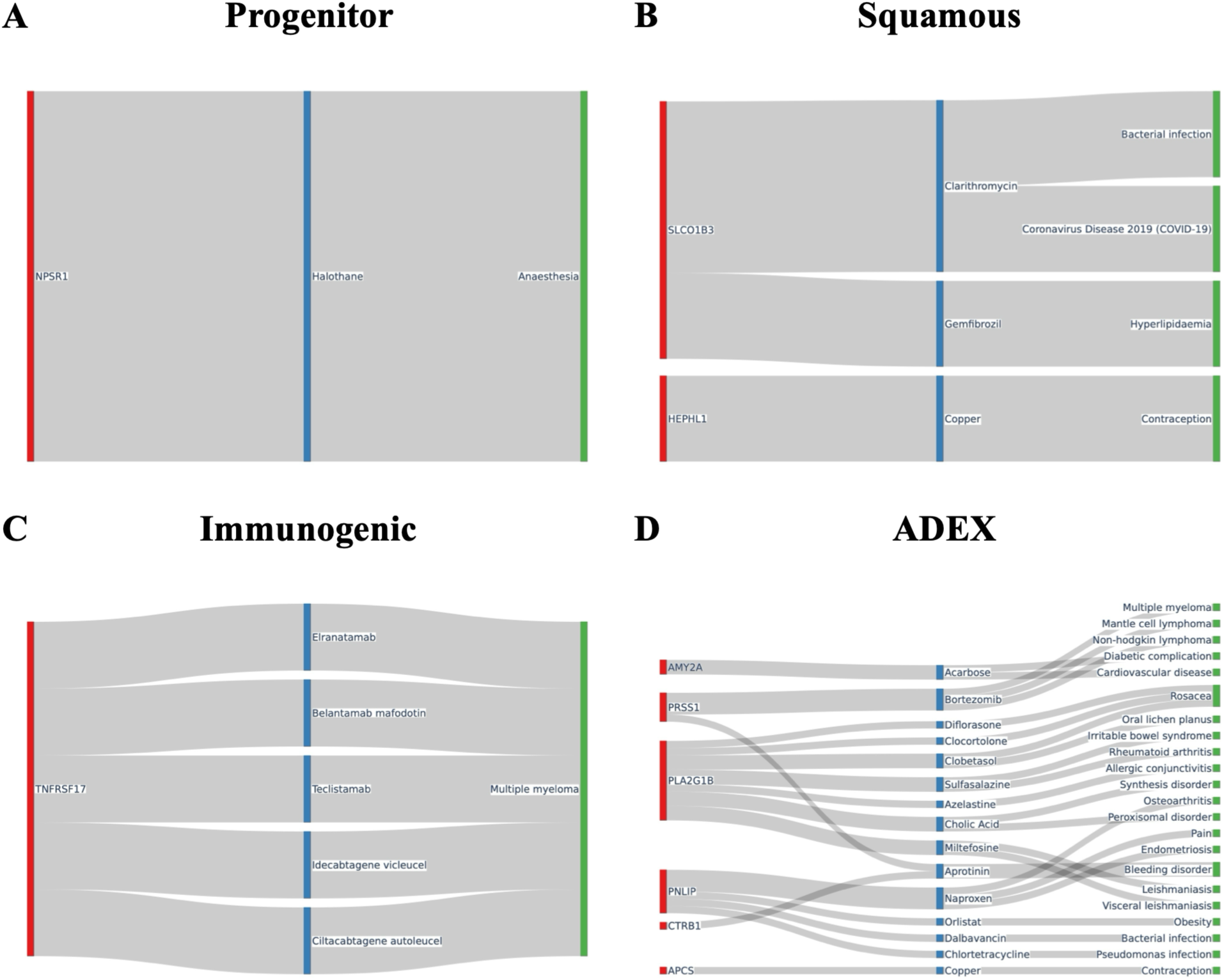
Drug and disease pathway associations for differentially expressed genes in each subtype. Sankey diagrams illustrating connections between highly differentially expressed genes and associated drug and disease pathways from the DrugBank database and Therapeutic Target Database (TTD) database for: (**A**) Progenitor, (**B**) Squamous, (**C**) Immunogenic, and (**D**) ADEX subtypes.

## Discussion

MetaPaCS is a novel stacking-based framework for predicting pancreatic cancer subtypes using RNA-seq data. Given the high complexity and heterogeneity of PaC transcriptomes, no single algorithm is universally suited for this classification task, and model performance is inevitably influenced by algorithmic bias. For example, linear SVM assumes linear separability and therefore cannot accurately estimate decision boundaries when the underlying relationships are non-linear^52^. Moreover, a comprehensive benchmark evaluating 179 algorithms across 121 datasets demonstrated that no individual model consistently outperformed others; instead, the top-performing algorithm varied substantially between datasets, underscoring the pervasive impact of algorithmic bias on classifier performance^53^. To mitigate these limitations, MetaPaCS integrated both stacking and meta-learning strategies. Stacking ensembles leverage the complementary strengths of diverse classifiers to generate a consensus prediction, while meta-learning enables the model to correct the systematic biases present in its base learners. The results suggest that our method improved the accuracy of the subtype predictions by around 1% in the best performing meta-learning classifier when compared to the best performing base classifier in the same run. Furthermore, concatenating the filtered expression data with the probability outputs from block A provides a richer, subtype-informed feature representation for block B, enabling the meta-learning classifier to simultaneously exploit the strengths of the base models and retain essential transcriptomic signals. As a result, MetaPaCS achieves performance comparable to or exceeding that of the best base classifier, while offering improved robustness across datasets. It was also observed that all the highest performing classifier combinations involve at least one SVM as the classifier. This is likely due to the higher performance of SVMs in cases where there is a relatively small sample number of high dimensional samples. From the different tests and measured performance of the meta-learner, we determined the optimal model for MetaPaCS to be all 10 base classifiers with linear SVM as the Meta-learner due to its consistently high performance.

In terms of visualization capabilities, MetaPaCS provided a more informative representation of pancreatic cancer transcriptomes than raw expression data. Conventional dimensionality-reduction methods applied directly to the original matrix produced substantial subtype overlap, reflecting the complexity of the data and the inability of unsupervised embeddings to recover subtype structure. In contrast, applying these methods to the MetaPaCS-derived embedding yielded markedly improved separation, indicating that the model captures discriminative signals not apparent in the raw features. Incorporating one-hot encoded subtype probabilities further enhanced cluster distinction, improving the interpretability of the visual space. These findings highlight the value of MetaPaCS in generating subtype-aware representations that enable clearer and more meaningful visualization of PaC heterogeneity. Differential gene expression analysis of the four subtypes also revealed a variety of important, uniquely upregulated genes that are essential to take into account when giving treatment to a patient with pancreatic cancer. For example, progenitor displays an upregulation in a gene responsible for the metabolic creation of glucose (G6PC2) and ADEX displays an upregulation in a gene responsible for the digestion of starch (AMY2A), which both serve to heighten blood sugar levels long term. This high blood sugar not only elevates the risk of type 2 diabetes but additionally may encourage cancer cell proliferation with the more easily available energy source. Another example is the upregulated genes in the squamous and immunogenic subtypes, which both influence the cancer characteristics in a way that makes treatment differ from progenitor or ADEX. Squamous, the most lethal subtype, shows many upregulated genes that are also found in other more lethal cancers such as MIR205HG, which is associated with migration and proliferation, and others like KRT6C, which are found to be upregulated in Lung adenocarcinoma. These genes likely influence the way a squamous subtype tumor would spread and metastasize, making it an important consideration for how treatment should be carried out. Immunogenic is similar, as the upregulated immunoglobin genes create an immunosuppressant microenvironment, which is important to take into consideration for immunotherapy-based treatments.

In addition, our GO pathway analysis found that each subtype showed upregulation in distinct pathways that lead to the unique characteristics of each subtype. The squamous subtype for example showed a high number of upregulated genes in pathways related to skin/dermis development and cellular structure, which would possibly result in a different tumor structure and different metastatic characteristics of tumors in this subtype. Additionally, 4 of the 15 most upregulated GO pathways in the squamous subtype were found to play a part in kidney development, possibly serving as a driving factor in some of the unique characteristics seen in this subtype. The immune subtype meanwhile shows upregulation in a multitude of immune response pathways, which was expected based on its characteristics and previous research. The Progenitor and ADEX subtypes are the most similar of the 4 subtypes, with both having upregulation in pathways meant mostly for the transport and localization of hormones and proteins. The main difference between the 2 is the upregulation in digestion and insulin secretion pathways seen in the Progenitor subtype.

Pathway analysis also revealed further medical considerations that should be taken for those with specific subtypes. One of the most notable being the upregulation of NPSR1, a gene linked to the metabolism of halothane, a common anesthetic used in developing countries. This upregulation may have unintended side effects on patients who use halothane, such as possibly increasing the known side effects like hepatotoxicity or cardiovascular disease due to the need for more anesthetic than the average patient. The ADEX subtype’s upregulated genes meanwhile effect 15 different medications, with the medications themselves showing being involved in the treatment of a variety of diseases ranging other cancers to diabetes and obesity. Without proper subtype diagnosis, the risk that these other diseases pose likely increases drastically due to overlooked changes in drug response.

While the results are promising, more research is needed to further optimize and evaluate MetaPaCS before it sees any major use in the medical field. Further research would include a far larger and more diverse dataset to obtain a more clear image of both the subtypes and the contribution the stacking aspect of the model makes to subtype prediction. In summary, MetaPaCS provides a robust and flexible framework for pancreatic cancer subtype classification by integrating diverse predictive signals into a unified, subtype-aware representation. We believe that MetaPaCS will significantly enhance downstream risk stratification for PaC and contribute to the design of more personalized treatment strategies.

## Conclusion

In this paper we introduced MetaPaCS, a high performing ensemble meta-learning method for the classification of pancreatic cancer into subtypes. Our method is based on a stacking ensemble, using the probability predictions of 10 base classifiers concatenated to the original input data as input for a new linear SVM meta-learning classifier. MetaPaCS performed significantly better than existing state-of-the-art methods for PaC subtyping and may be able to have a positive impact on PaC diagnosis and treatment in the future, allowing for safer and more personalized treatment for PaC patients while taking the potential additional health considerations into effect, drastically lowering the mortality rate. Future research aims to make use of a much larger pancreatic cancer dataset with the inclusion of multi-omics data integration. Given adequate data, this model’s framework may be extensible to subtype prediction in other cancer subtypes as well.

## Author contributions

The idea for this study was conceived of and designed by SW. SW, MS, and NP developed the method. NP performed experiments and analysis on the data. All authors participated in the writing and revision of the paper. The manuscript was approved by all authors.

## Data Availability

All data used in this study are publicly available in the corresponding references mentioned in the manuscript.

## Code Availability

All code used for MetaPaCS is publicly available and can be found on GitHub at https://github.com/wan-mlab/MetaPACS.

## Supporting information

supplemental figures and their descriptions are in this powerpoint

