## supplemental figures and their descriptions are in this powerpoint for "MetaPaCS: A Novel Meta-Learning Framework for Pancreatic Cancer Subtype Identification"

### Slide 1
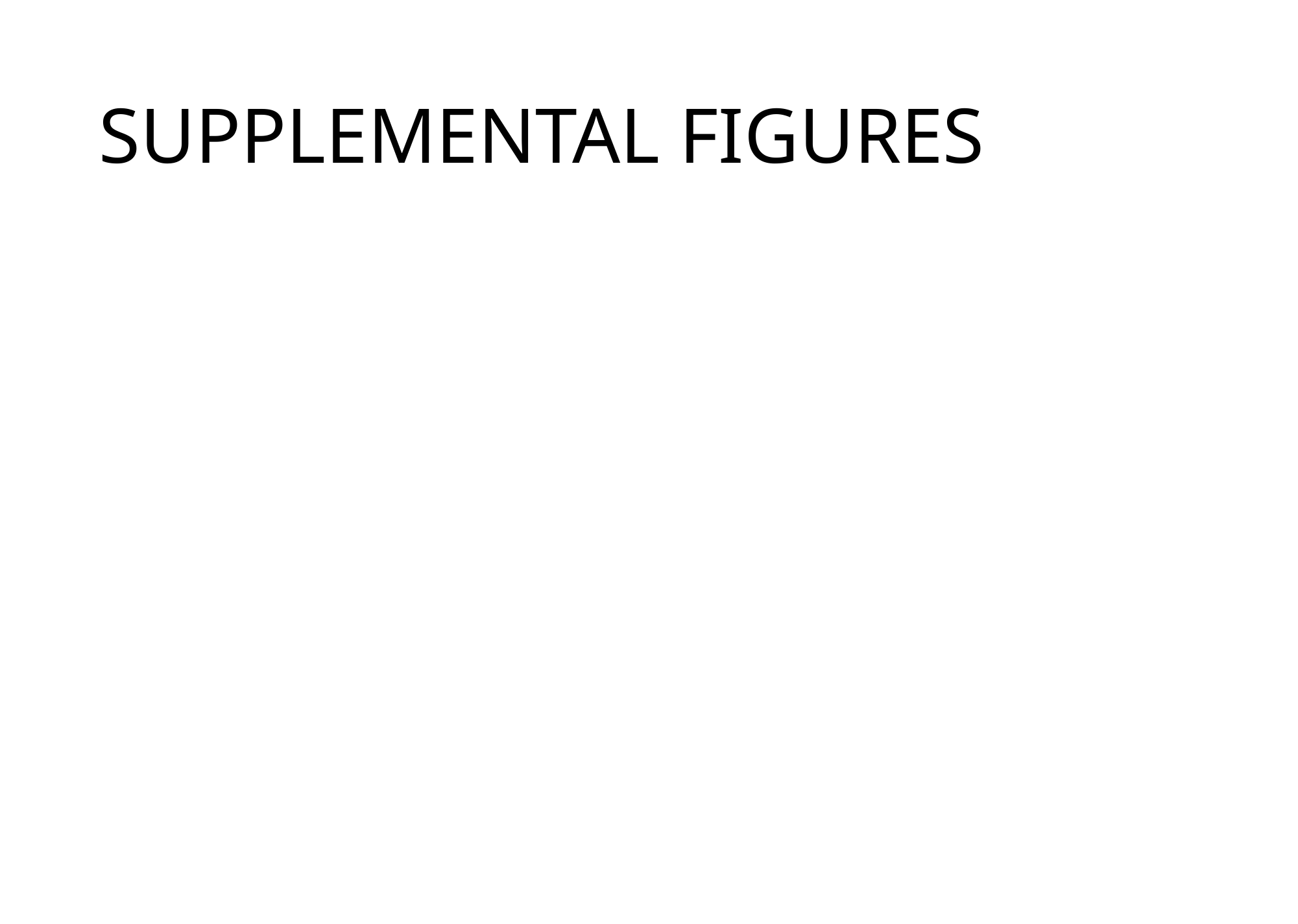

#### SUPPLEMENTAL FIGURES

### Slide 2
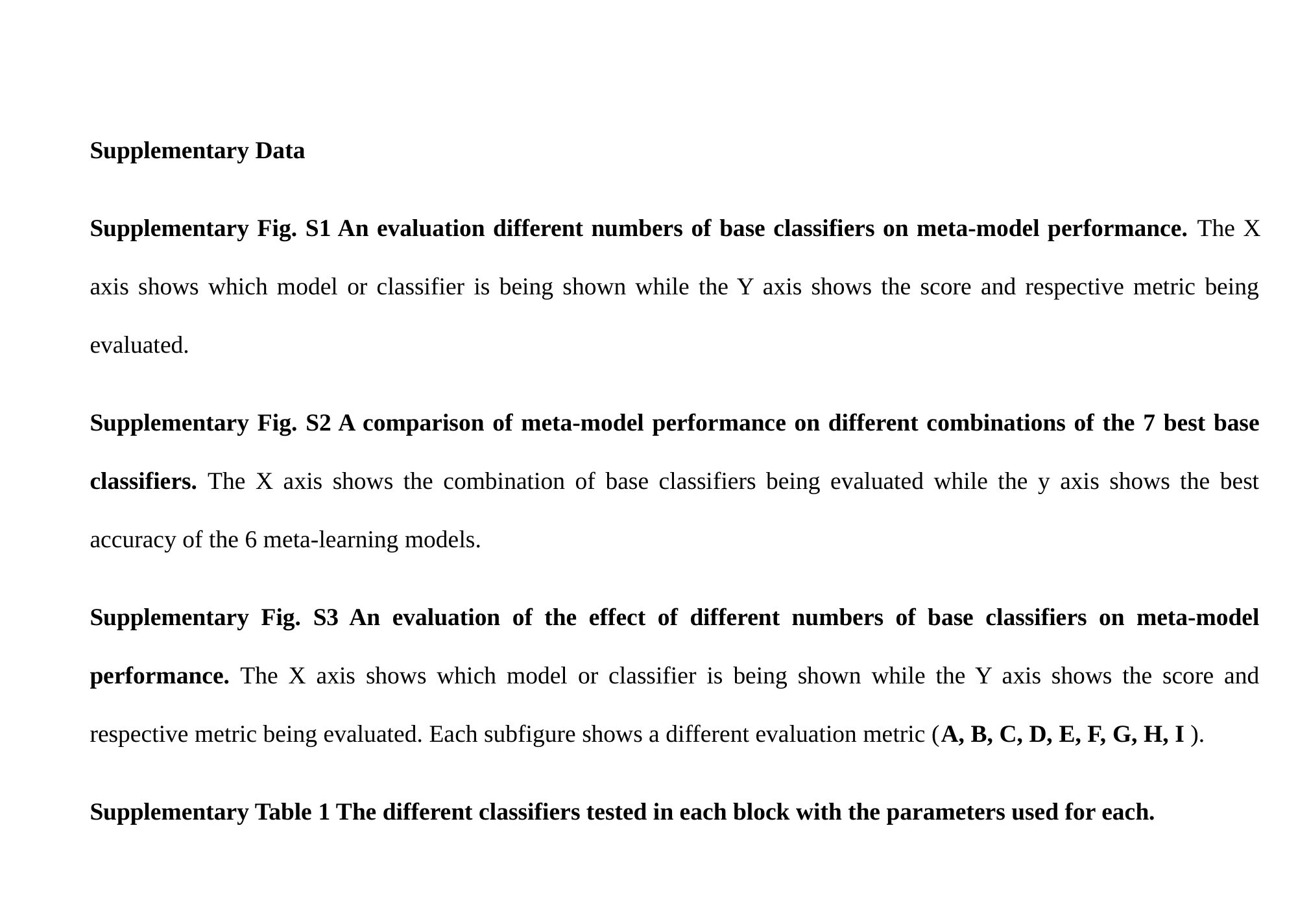

Supplementary Data
Supplementary Fig. S1 An evaluation different numbers of base classifiers on meta-model performance. The X axis shows which model or classifier is being shown while the Y axis shows the score and respective metric being evaluated.
Supplementary Fig. S2 A comparison of meta-model performance on different combinations of the 7 best base classifiers. The X axis shows the combination of base classifiers being evaluated while the y axis shows the best accuracy of the 6 meta-learning models.
Supplementary Fig. S3 An evaluation of the effect of different numbers of base classifiers on meta-model performance. The X axis shows which model or classifier is being shown while the Y axis shows the score and respective metric being evaluated. Each subfigure shows a different evaluation metric (A, B, C, D, E, F, G, H, I ).
Supplementary Table 1 The different classifiers tested in each block with the parameters used for each.

### Slide 3
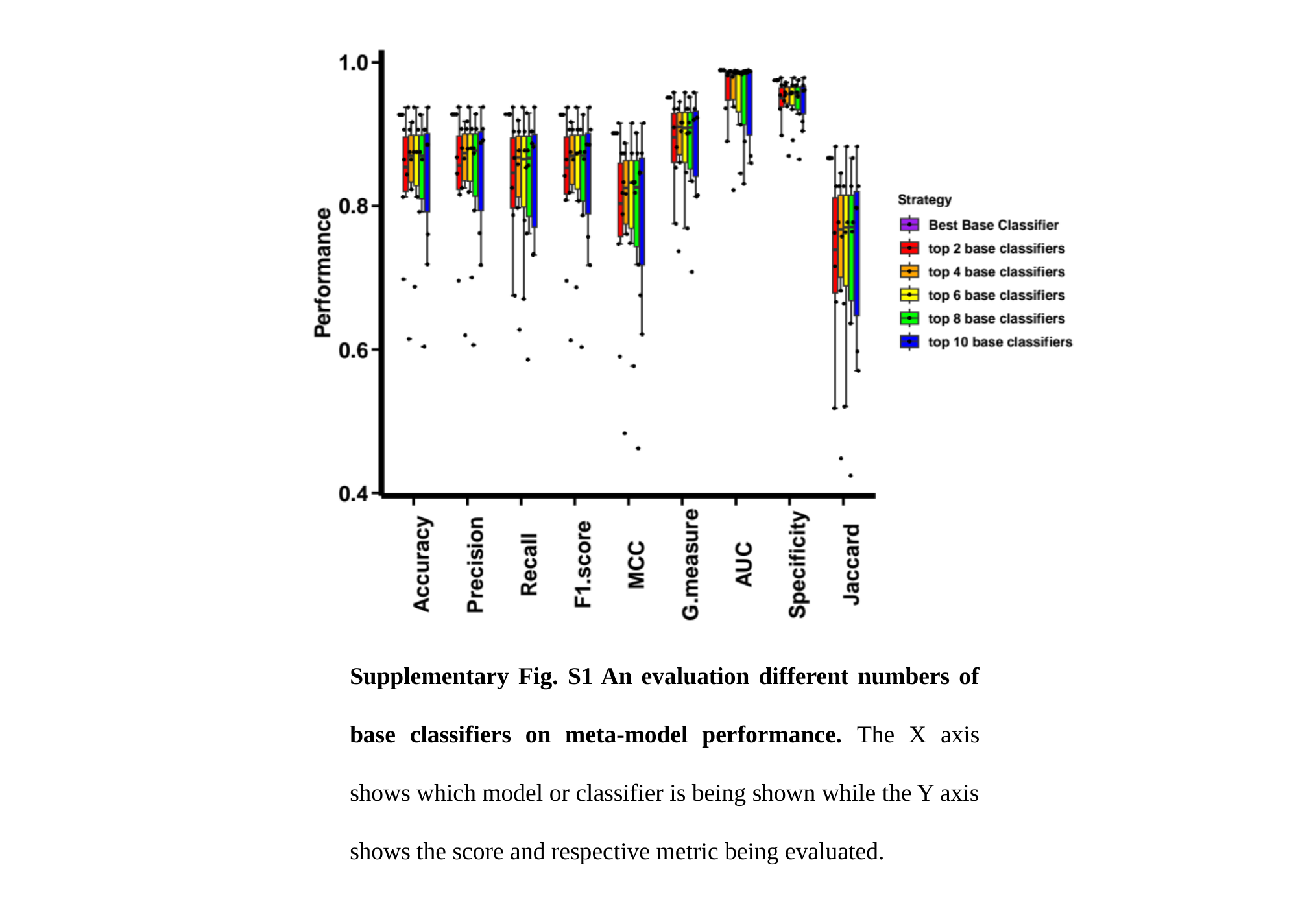

Supplementary Fig. S1 An evaluation different numbers of base classifiers on meta-model performance. The X axis shows which model or classifier is being shown while the Y axis shows the score and respective metric being evaluated.

### Slide 4
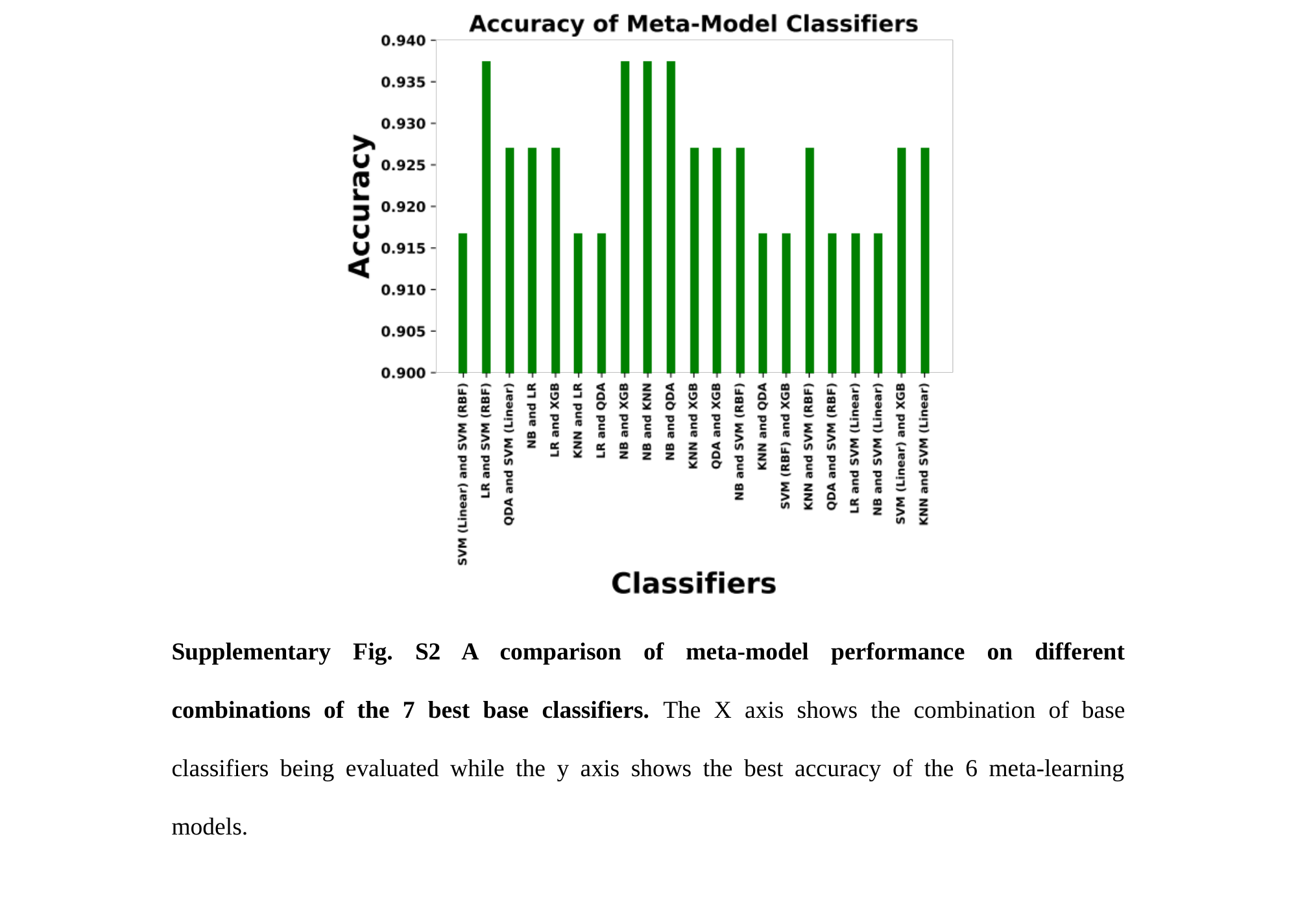

Supplementary Fig. S2 A comparison of meta-model performance on different combinations of the 7 best base classifiers. The X axis shows the combination of base classifiers being evaluated while the y axis shows the best accuracy of the 6 meta-learning models.

### Slide 5
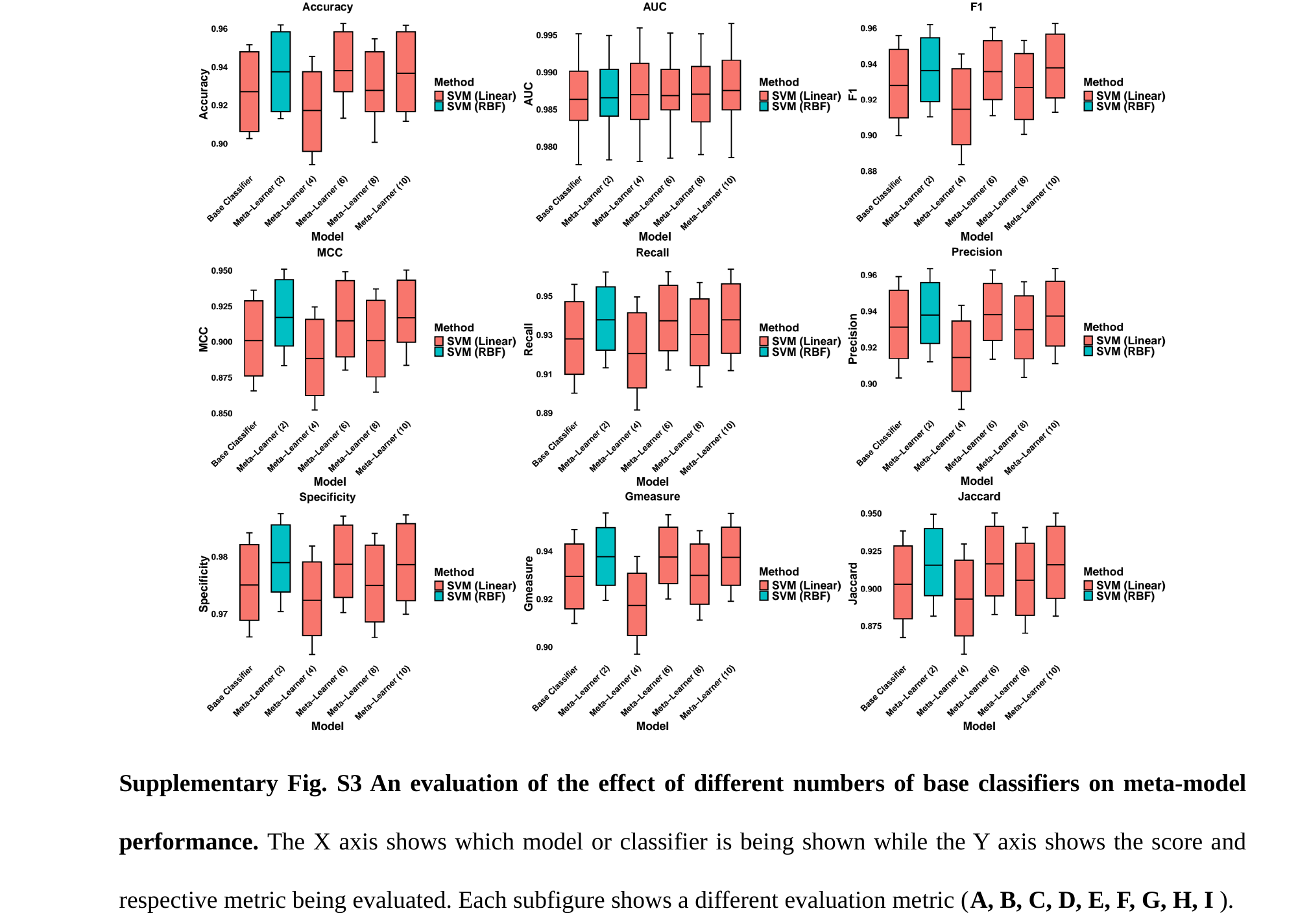

Supplementary Fig. S3 An evaluation of the effect of different numbers of base classifiers on meta-model performance. The X axis shows which model or classifier is being shown while the Y axis shows the score and respective metric being evaluated. Each subfigure shows a different evaluation metric (A, B, C, D, E, F, G, H, I ).

### Slide 6
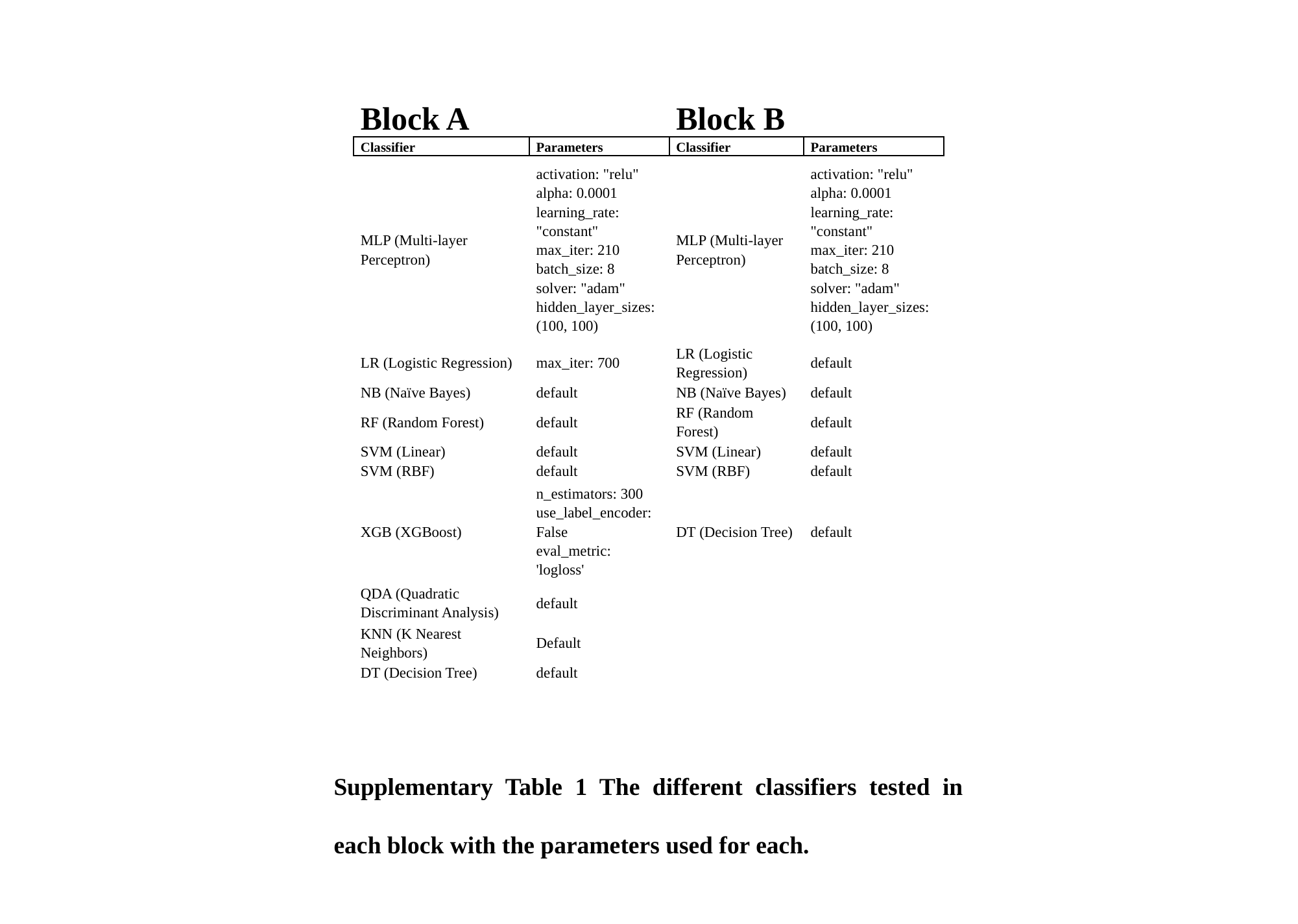

| Block A | | Block B | |
| --- | --- | --- | --- |
| Classifier | Parameters | Classifier | Parameters |
| MLP (Multi-layer Perceptron) | activation: "relu" alpha: 0.0001 learning\_rate: "constant" max\_iter: 210 batch\_size: 8 solver: "adam" hidden\_layer\_sizes: (100, 100) | MLP (Multi-layer Perceptron) | activation: "relu" alpha: 0.0001 learning\_rate: "constant" max\_iter: 210 batch\_size: 8 solver: "adam" hidden\_layer\_sizes: (100, 100) |
| LR (Logistic Regression) | max\_iter: 700 | LR (Logistic Regression) | default |
| NB (Naïve Bayes) | default | NB (Naïve Bayes) | default |
| RF (Random Forest) | default | RF (Random Forest) | default |
| SVM (Linear) | default | SVM (Linear) | default |
| SVM (RBF) | default | SVM (RBF) | default |
| XGB (XGBoost) | n\_estimators: 300 use\_label\_encoder: False eval\_metric: 'logloss' | DT (Decision Tree) | default |
| QDA (Quadratic Discriminant Analysis) | default | | |
| KNN (K Nearest Neighbors) | Default | | |
| DT (Decision Tree) | default | | |
Supplementary Table 1 The different classifiers tested in each block with the parameters used for each.
